# scMetaIntegrator: a meta-analysis approach to paired single-cell differential expression analysis

**DOI:** 10.1101/2025.06.04.657898

**Authors:** Kalani Ratnasiri, Sara N. Mach, Catherine A. Blish, Purvesh Khatri

## Abstract

Traditional differential gene expression methods are limited for analysis of single cell RNA-sequencing (scRNA-seq) studies that use paired repeated measures and matched cohort designs. Many existing approaches consider cells as independent samples, leading to high false positive rates while ignoring inherent sampling structures. Although pseudobulk methods address this, they ignore intra-sample expression variability and have higher false negatives rates. We propose a novel meta-analysis approach that accounts for biological replicates and cell variability in paired scRNA-seq data. Using both real and synthetic datasets, we show that our method, single-cell MetaIntegrator (https://github.com/Khatri-Lab/scMetaIntegrator), provides robust effect size estimates and reproducible p-values.

## Background

The ability to accurately detect and model individual RNA transcripts is integral to scRNA-seq methods. Advances in methods include modeling counts via negative binomial distributions or more complex methods such as MAST to account for undetectable and non-zero genes [1]. However, inherent complexities in gene expression profiling pose significant challenges as gene capture and expression measurements are influenced by a myriad of factors, including dropout events and variability of sequencing depth [2]. Both biological variations and technical artifacts can impact gene measurements, making it difficult to discern which factors are driving the observed results. These nuances lead to diverse distributions of gene expression, complicating the effective development of one-size-fits-all models to build differential gene expression methods and account for these inherent variations [3].

Equally important is the need to account for biological replicates when analyzing scRNA-seq data. Several single-cell methods, including the widely popular Wilcoxon rank-sum test, treat each individual cell as an independent sample, ignoring the cell’s sample of origin to thereby not accounting for biological correlation. This oversight often results in elevated false discovery rates (FDR) and inflated p-values, thus compromising the reliability of the findings [3–6]. Methods based on pseudobulk gene expression address this issue by aggregating gene expression values across all cells within a sample [7,8]; however, these methods fail to account for the intrinsic cell-to-cell variability in gene expression and tend to have higher false negative rates [8]. Applications of mixed-model approaches to effectively account for paired sampling are hampered by lengthy computational times that do not consistently yield superior results compared to pseudobulk and Wilcoxon rank-sum methods [4].

Further, more complex scRNA-seq study cohort designs that include paired sampling, such as matched and longitudinal cohorts, require analysis using methods that account for both sample of origin as well as the inherent paired structure design. While many pseudobulk methods can account for additional sampling structure, few single-cell methods can take these structures into account. This underscores the need for more robust and easy-to-use analytical techniques to enable users to analyze their scRNA-seq data appropriately.

In response to the complex challenges associated with scRNA-seq data analysis, we developed scMetaItegrator, which leverages the meta-analysis framework to identify differentially expressed genes in paired scRNA-seq samples. It is built upon our previously published MetaIntegrator method [9,10] that was originally designed for bulk transcriptomic data to identify robust and reproducible DEGs and disease signatures [11–13]. scMetaIntegrator extends these capabilities to paired scRNA-seq cohorts, including matched and longitudinal sampling. Our results show scMetaIntegrator has higher accuracy compared to popular pseudobulk and single-cell differential expression (DE) approaches for identifying consistently DEGs across sample pairs and calculating robust magnitudes of change values. Moreover, it has higher sensitivity for identification of DEG compared to pseudobulk methods, and higher specificity as compared to other single-cell DE methods. Our analysis also found that scMetaIntegrator can identify DEGs with consistent p-values and effect sizes even with reduced sample sizes in both real-world and simulated data. To facilitate its widespread use, we have developed an R package that integrates seamlessly with the existing MetaIntegrator framework and Seurat pipeline, allowing ease of adoption while integrating with MetaIntegrator’s suite of visualization tools to advance gene expression analysis and generate meaningful biological insights from complex scRNA-seq data.

## Results

### Single-cell meta-analysis methods account for biological replicates and cell variability

Current single-cell DE methods fall in two broad categories: single-cell and pseudobulk methods. Single-cell DE methods typically consider each cell as an independent measurement, which can lead to high false positive rates (**Fig. 1A**). Alternatively, psudobulk DE methods rely on creating “pseudobulk” transcriptome profiles that consider all the cells from an individual as a single measurement and tend to have reduced sensitivity (**Fig. 1B**) [8]. To improve upon both classes of methods, we developed scMetaIntegrator, which accounts for both the biological sample and the inherent gene expression variability across its cells. We leverage a meta-analysis framework for a scRNA-seq study by considering each paired sample as a distinct “dataset,” with individual cells from each pair serving as the “samples” for each dataset (**Fig. 1C**). For each gene, we calculate an effect size as Hedges’ *g*, which accounts for the mean gene expression across each pair of samples and the pooled standard deviation of expression values of all cells within each dataset. Finally, we use a random effects inverse variance model to estimate summary effect sizes under the assumption that the true change in gene expression across groups may vary across different pairs. This assumption further accounts for the inherent variability across pairs that may be driven by matched features.

**Figure 1.**
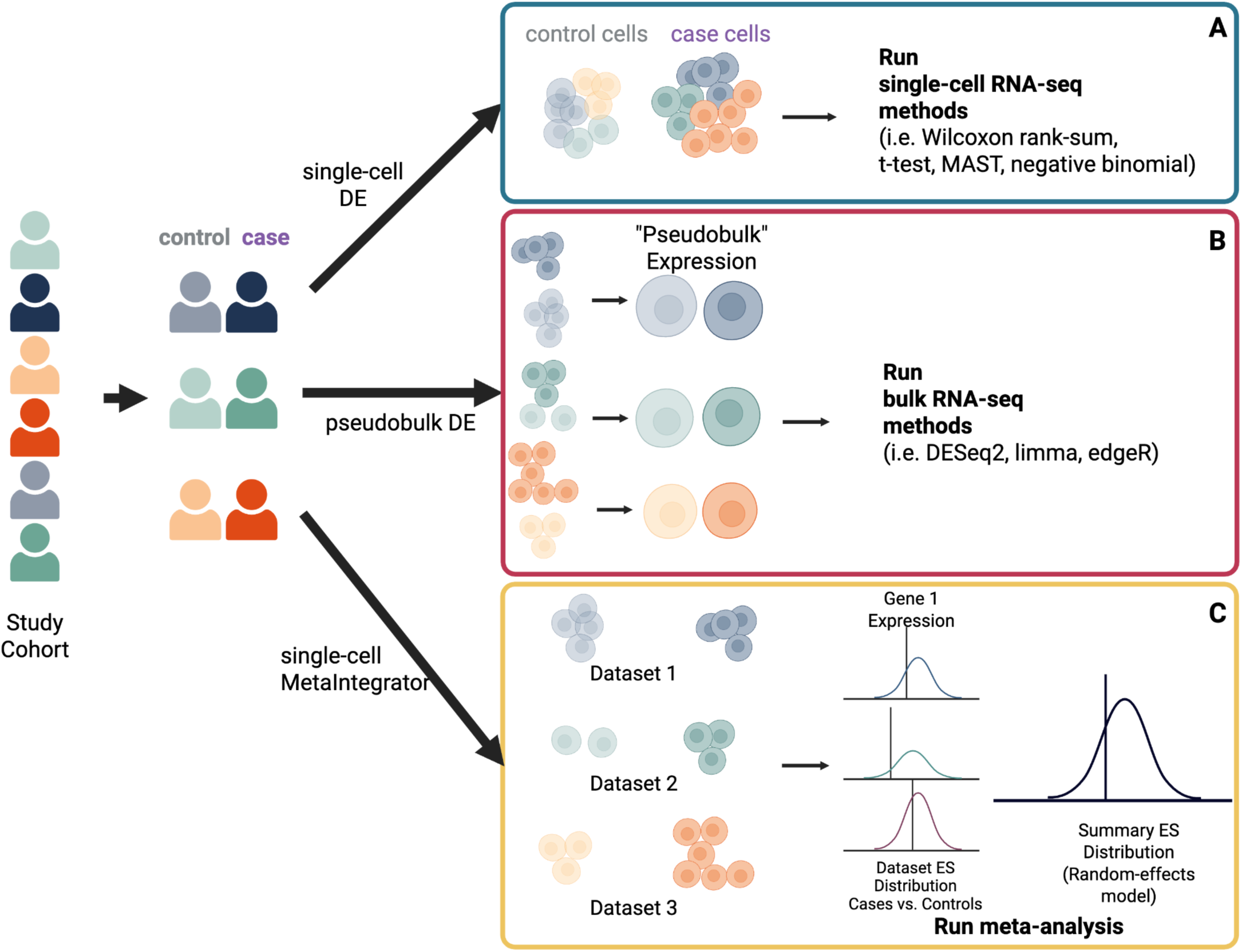
Schematic of single cell differential expression methods. **(A-C)** Schematic for **A)** single-cell DE processing, **B)** pseudobulk DE processing, and **C)** single-cell MetaIntegrator processing.

### Comparing scMetaIntegrator to single-cell and pseudobulk DE methods across matched cohort data

To investigate the ability of scMetaIntegrator to account for matched cohort designs, we used a scRNA-seq dataset of peripheral blood mononuclear cells (PBMCs) from individuals with celiac disease and matched healthy controls (**Fig. 2A**) [14], which included eight pairs, each composed of a healthy control and an individual with celiac disease matched for age-, sex-, and human leukocyte antigen (HLA)-genotype. To limit any cell-annotation bias, throughout our manuscript, we analyze single-cell data as complete samples, without separating cell types.

**Figure 2.**
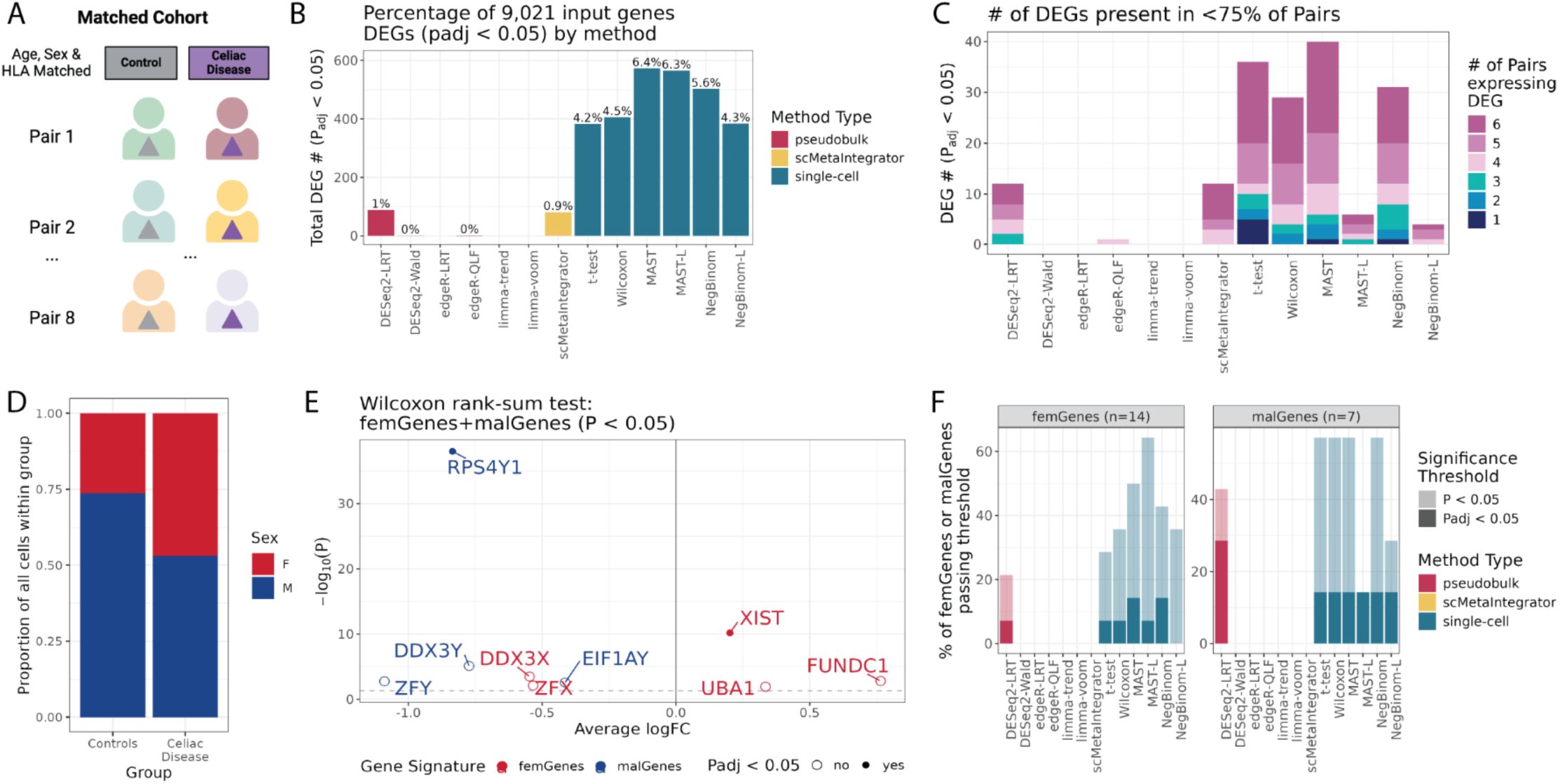
Single-cell MetaIntegrator identifies conserved DEGs across paired comparisons while accounting for matched features. **A)** Schematic illustration of the public dataset from Ramírez-Sánchez et al. used for the entirety of the figure. **B)** Of the 9,021 genes that were expressed across at least 1% of cells within either group, the number of significant genes identified by each DE method (padj < 0.05). **C)** Stacked barplot of the subset of DEGs from **B** that were expressed in at least 1% of cells of only 1 - 6 sample pairs. **D)** The proportion of cells in each group that are from either male or female donors. **E)** Genes identified by Wilcoxon rank-sum DE analysis at p < 0.05 or padj < 0.05 that are part of the femGene or malGene gene signature. Average log fold-change (logFC) calculated through Seurat analysis. **F)** Overlay barplot of the percentage of genes from the femGenes or malGenes gene signatures that are identified by each DE method that pass either p < 0.05 or padj < 0.05 thresholds. In each facet header is the number of genes from each signature represented across the 9,021 thresholded genes prior to DE analysis.

We performed DE analyses using scMetaIntegrator, 6 pseudobulk methods (DESeq2-LRT, DESeq2-Wald, edgeR-LRT, edgeR-QLF, limma-trend, limma-voom) and 4 single-cell methods (t-test, Wilcoxon rank-sum, MAST, negative binomial test) [4]. All pseudobulk methods and two of the single-cell methods accounted for pairing between samples. For single-cell methods “-L” denotes the use of pairing as a latent variable in the method’s model. Single-cell methods consistently identified a higher number of DEGs (4.2-6.4%) compared to pseudobulk (<1%) and scMetaIntegrator (0.9%), despite using more stringent p-value adjustment (Bonferroni correction vs. FDR; **Fig. 2B**).

We hypothesized that the high number of DEGs identified by single-cell methods may be driven by individual samples in the methods that do account for the sample of origin. Therefore, we evaluated the consistency of the expression of these DEGs across pairs by quantifying the number of sample pairs that expressed DEGs for each method in at least 1% of all cells. We found that 5,994 (66%) of the 9,021 genes included in the analysis were consistently expressed across 7-8 pairs. However, several DEGs identified by methods were expressed in fewer pairs, with single-cell methods identifying substantial number of genes (up to 10%) expressed in half or less than half of the pairs (**Fig. 2C**). Single-cell methods, which accounted for sample pairs, identified a lower number of inconsistently expressed DEGs, suggesting that including pairing in models for MAST and the negative binomial test may reduce sample bias in DEG identification. These results highlighted that methods that do not account for samples identify DEGs that may be changing inconsistently across the groups of interest and be driven by outlier samples.

Next, we investigated whether despite matching, the methods that identified genes with inconsistent expression across pairs as significant were capturing variations related to unequal sample representation. The proportion of female cells was higher across the celiac disease group than the healthy control group (**Fig. 2D**). Therefore, we hypothesized that this difference would be identified by methods that do not account for the pairing structure. To test this hypothesis, we investigated whether each method identified sex-related genes as significant using 21 genes from two published sex signature gene sets of 14 X-escape genes ("femGenes") and 7 Y-chromosome genes ("malGenes")[15]. Consistent with the differences in cell proportions, method using Wilcoxon rank-sum test found 9 out of 21 sex-specific genes as significantly different such that multiple female genes were over- expressed in celiac patients, whereas 4 out of 7 male genes were over-expressed in healthy controls (**Fig. 2E**). Similarly, all single-cell methods identified at least 10% of sex-related genes as statistically significant, where scMetaIntegrator did not identify any sex-related genes as significant (**Fig. 2F**). These results suggest that the false positive rate for single-cell methods is substantially higher.

Together, these results demonstrate that scMetaIntegrator, unlike other single-cell methods, is more robust to cell and gene measurement imbalances across groups by accounting for pairing, which in turn leads to the identification of DEGs consistently expressed across matched pairs, substantially reducing false positives potentially arising from variations in sample representation.

### Comparing scMetaIntegrator to single-cell and pseudobulk DE methods across repeated measures data

We compared scMetaIntegrator to other methods for the analysis of longitudinal samples obtained between groups with strong transcriptional changes using a longitudinal scRNA-seq dataset of PBMCs from donors pre- and post-vaccination [16]. We used PBMCs from five donors at pre- and one day post- vaccine boost – timepoints previously shown to exhibit pronounced transcriptional changes (**Fig. 3A**). Here, we used 10,374 genes expressed in at least 1% of cells within each sample pair to focus on changes in genes that are consistently expressed in all pairs.

**Figure 3:**
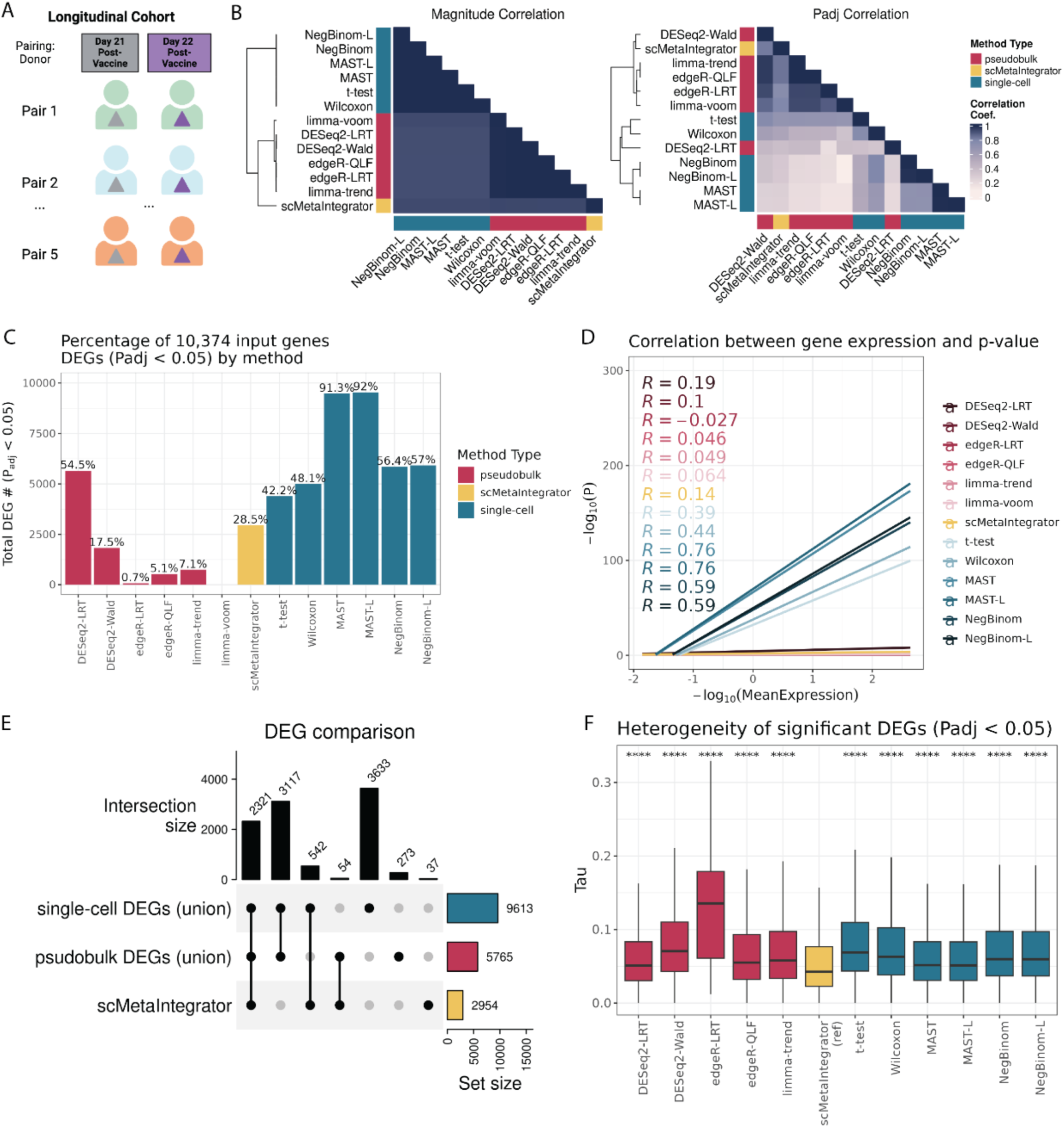
Single-cell MetaIntegrator identifies consistently changing DEGs across paired comparisons. **A)** Schematic illustration of the Arunachalam et al. public dataset used across the figure. **B)** Correlation plots of the Spearman’s rank correlation coefficients of comparisons of the 10,374 genes that were expressed across all 5 donor pairs in at least 1% of cells within either pairing by the (left) magnitude of change and (right) adjusted p-value values between each method. **C)** The number of significant DEGs identified by each DE method (padj < 0.05). **D)** Spearman’s rank correlation of the p-values compared to the overall average expression of the gene across all cells in the dataset. **E)** Upset plot comparison of all significant DEGs (padj <0.05) identified by single-cell methods, pseudobulk methods, and scMetaIntegrator. **F)** Boxplot of the tau values of all significant DEGs (padj < 0.05) identified by each method. Bonferonni adjusted one-sided t-test was performed, testing whether each method’s DEGs had tau values greater than those from DEGs identified by scMetaIntegrator. Asterisk values across figures are represented as follows: ∗padj value < 0.05, ∗∗∗padj value < 0.01, ∗∗∗∗padj value < 0.001, and ∗∗∗∗∗padj value < 0.0001.

To characterize the relationship between different DE methods, we performed Spearman’s rank correlation analysis comparing both magnitude of change and adjusted p-values for each pairwise combination of methods. First, methods of similar type (e.g., single-cell or pseudobulk) were more correlated with each other, where scMetaIntegrator correlated more with pseudobulk methods (**Fig. 3B**). When correlating adjusted p-values, the correlations between methods varied widely (0.09<=r<=1) compared to those for the magnitude of change (0.93<=r<=1). Furthermore, each of the single-cell DE methods identified a large percentage of the 10,374 measured genes as DEGs, ranging from 42.2% (4378 genes) for t-test to 92% (9544 genes) for MAST-L (**Fig. 3C**). In contrast, all pseudobulk methods, except DESeq2-LRT, identified a substantially lower number of DEGs, ranging from 0.7% (73 genes) for edgeR-LRT to 17.5% (1816 genes) for DESeq2-Wald. Compared to the single-cell and pseudobulk methods, scMetaIntegrator identified 28.48% of DEGs. This result suggests that although most methods estimate expression changes similarly, they vary substantially in how they identify DEGs.

Previous studies have found that statistical significance (i.e., p-values) is strongly correlated with expression level in single-cell DE analysis [4,17,18], which leads to bias towards highly expressed genes being identified as DEGs. Therefore, we investigated whether statistical significance in scMetaIntegrator is independent of expression. In line with previous studies, other single-cell methods had significant correlations between p-values and gene expression (0.39<=r<=0.76). In contrast, scMetaIntegrator had substantially lower correlation (r=0.14), comparable with the weak correlations observed with most pseudobulk methods (-0.027<=r<=0.19; **Fig. 3D**). Thus, scMetaIntegrator and pseudobulk methods are less likely to identify a gene as significantly different based on high expression levels.

When we compared significant DEGs (i.e., those with padj<0.05) between the individual methods using Jaccard Index, these methods broadly clustered by single-cell and pseudobulk methods (**Fig. S1A**). While single-cell methods identified thousands of different DEGs, pseudobulk methods and scMetaIntegrator still identified unique DEGs (**Fig. 3E**). For example, *CTNNAL1,* gene identified as the most significant only by scMetaIntegrator, was consistently upregulated across all pairs (**Fig. S1B-C**). This result demonstrates that despite other methods identifying thousands of significant genes, many true positives are still missed. On the other hand, *ZBTB1*, a gene identified as significantly upregulated by all single-cell methods, increased in only one donor at day 22 (**Fig. S1E-F**), indicating that it is likely a false positive due to an outlier sample. Similarly, *CD226*, a gene significant across four out of six pseudobulk methods (**Fig. S1G**), had inconsistent changes across pseudobulk results (**Fig. S1H-I**).

Because of the between-pair variability of DEGs identified by specific examples (**Fig. S1**), we hypothesized that scMetaIntegrator identified DEGs with consistent expression changes across sample pairs and is more resistant to outlier samples. To test this, we compared the heterogeneity in effect sizes between sample pairs, measured as tau for each gene. Heterogeneity between sample pairs for DEGs identified by all methods were significantly higher compared to that of DEGs identified scMetaIntegrator (padj<0.05, one-sided t-test) (**Fig. 3F**). In other words, DEGs identified by scMetaIntegrator had consistent changes across sample pairs than the DEGs identified by other methods. When comparing the standard deviation of paired logFC generated from the aggregated pseudobulk data (**Fig. S2A**), scMetaIntegrator had the lowest standard deviation than other methods, further demonstrating that scMetaIntegrator identifies consistently changing DEGs.

To further understand the effect of outliers on identification of DEGs, we compared the direction of change associated with each method and determined which of those changes were consistent with only a single donor’s direction of change (**Fig. S2B**). All single-cell methods, except scMetaIntegrator, identified up to 9.8% of DEGs where the magnitude of change was consistent with the change of only one of the five donors. On the other hand, all pseudobulk methods, except DESeq2-LRT, and scMetaIntegrator were resistant to outliers.

These results provide empirical evidence that scMetaIntegrator is resistant to potential biases introduced by outlier samples and individual sample variations. Avoiding these biases is crucial for identifying genes that are different between groups and robust to spurious signals arising from inter-individual variability.

### scMetaIntegrator has lower false discovery rates than single-cell methods in simulation data

We further investigated the robustness of scMetaIntegrator to outlier samples using simulation data with known ground truth changes. We compared scMetaIntegrator with the three most-used single-cell DE methods (DESeq2, Wilcox, and MAST) [4].

We used ‘splatter’ [19,20] to generate a dataset composed of 4 donors with samples divided between two groups. We simulated this data to have a set of DEGs with known magnitude of change. Next, we simulated Donor1 as an outlier sample with an underlying condition driving gene expression differences from the other 3 donors (**Fig. 4A**). To determine how the outlier donor impacted DE results, we used 1000 cells from all samples from donors 2-4, while varying the number of cells from the outlier Donor1 from 50 to 5000 cells per sample.

**Figure 4:**
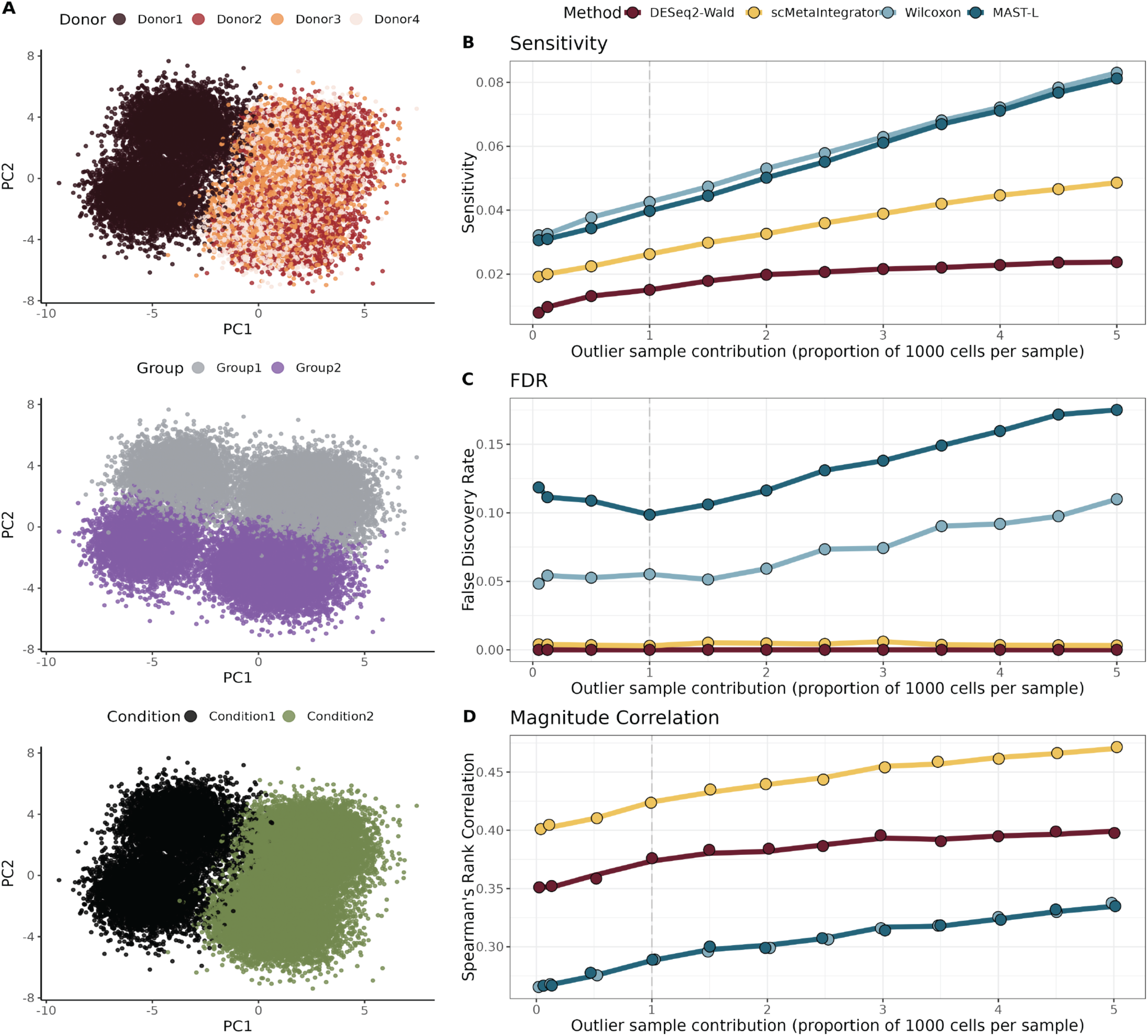
Comparative DE analysis across simulated outlier single-cell data. **A)** PCA plots of the original simulated data, composed of four donors each evenly split into two samples, 5000 cells each, between two groups. Donor1 is the outlier donor, simulated with an underlying condition. **(B-D)** Values of power, FDR, and magnitude of change correlation measurements as each DE method is run on a dataset where the proportion of Donor1’s cells varies. A random subset of 1000 cells/sample were included in analysis for Donors 2-5. For Donor1, the proportion of cells ranged from 5% (50 cells/sample) to 500% (5000 cells/sample) of the number of cells from the other donors. Dashed line at x = 1 represents when all samples are at equal cell numbers. **B)** Discovery power (TP/ (TP+FN)) of each method’s DEGs (padj <0.05) as the contribution of outlier sample cells varies. **C)** False discovery rate (FP/ (FP+TP)) each method’s DEGs (padj <0.05) as the contribution of outlier sample cells varies. **D)** Spearman’s rank correlation of each method’s DE magnitude of change estimates compared to the simulated changes as the contribution of outlier sample cells varies.

We first tested the discovery power of each of the methods to identify true DEGs (Fig. 4B). Across all simulations, Wilcoxon and MAST-L had the highest sensitivity, DESeq2-Wald had the lowest sensitivity, and scMetaIntegrator had consistently higher sensitivity compared to DESeq2-Wald. Sensitivity for the single-cell methods and scMetaIntegrator increased linearly with increased number of cells from Donor1.

Next, we measured the false discovery rate (FDR) within DEGs identified by each method as the number of cells from the outlier donor increased (**Fig. 4C**). Irrespective of the number of cells used, FDRs for Wilcoxon and MAST-L were more than ten times those of scMetaIntegrator. Notably, the FDRs of Wilcoxon and MAST-L were consistently higher than their sensitivity rates, irrespective of the number of cells from the outlier sample. In other words, although Wilcoxon and MAST-L had higher sensitivity at the cost of substantially higher FDRs. Further, FDRs for Wilcoxon and MAST-L increased linearly as the number of cells from the outlier donor samples increased, whereas FDRs for scMetaIntegrator and DEseq2-Wald remained unchanged. Together, these results suggest that the increased rate of true DEGs in Wilcoxon and MAST-L is accompanied by even higher levels of false positives, highlighting an unfavorable tradeoff. In contrast, scMetaIntegrator demonstrated higher sensitivity that comes with increased sample size, while also substantially reducing false positives.

Finally, we tested how magnitude of change estimates correlated with the true simulated changes in the genes across groups for each method (**Fig. 4D**). Across all methods, irrespective of the number of cells from the outlier donor samples used, the effect size estimates by scMetaIntegrator had highest correlation with the true DEG changes than the other methods. The single-cell methods had the least correlation. When we repeated the same analyses such that there was no outlier donor, but had variability in overall inter-individual similarity, we found similar patterns across methods such that scMetaIntegrator produced the most correlated effect size estimates than other methods (**Fig. S3**).

Collectively, our results demonstrate that scMetaIntegrator has higher power to identify true DEGs than pseudobulk-based methods (e.g., DESeq2-Wald), lower FDRs than single-cell methods (e.g., Wilcoxon, MAST- L), and more accurate estimates changes in expression across all methods, despite an outlier donor.

### Comparing the biological reproducibility of scMetaIntegrator to other DE methods

We next examined reproducibility of DE results across methods. We used a dataset by Kang *et al.* (referred to as kang2017 dataset) [21] that profiled PBMCs from 8 donors with and without interferon-beta (IFN-ꞵ) stimulation using scRNA-seq (**Fig. 5A**). As validation data, we identified an independent scRNA-seq data generated from Parse Biosciences that profiled PBMCs from 12 donors with and without IFN-ꞵ-stimulation using scRNA-seq (**Fig. 5A**) [22]. We compared these methods using three metrics: 1) area under the concordance curve (AUCC), which compares the order of genes by multiple hypotheses corrected p-values between two comparisons 2) fold- change concordance (FCC), where adjusted p-values are multiplied by the sign of the magnitude of change and these results values were compared via Spearman’s rank correlation; 3) Pearson correlation of magnitude of change estimates. FCC here highlights the combination of results from both p-value estimates and magnitude of change estimates.

**Figure 5:**
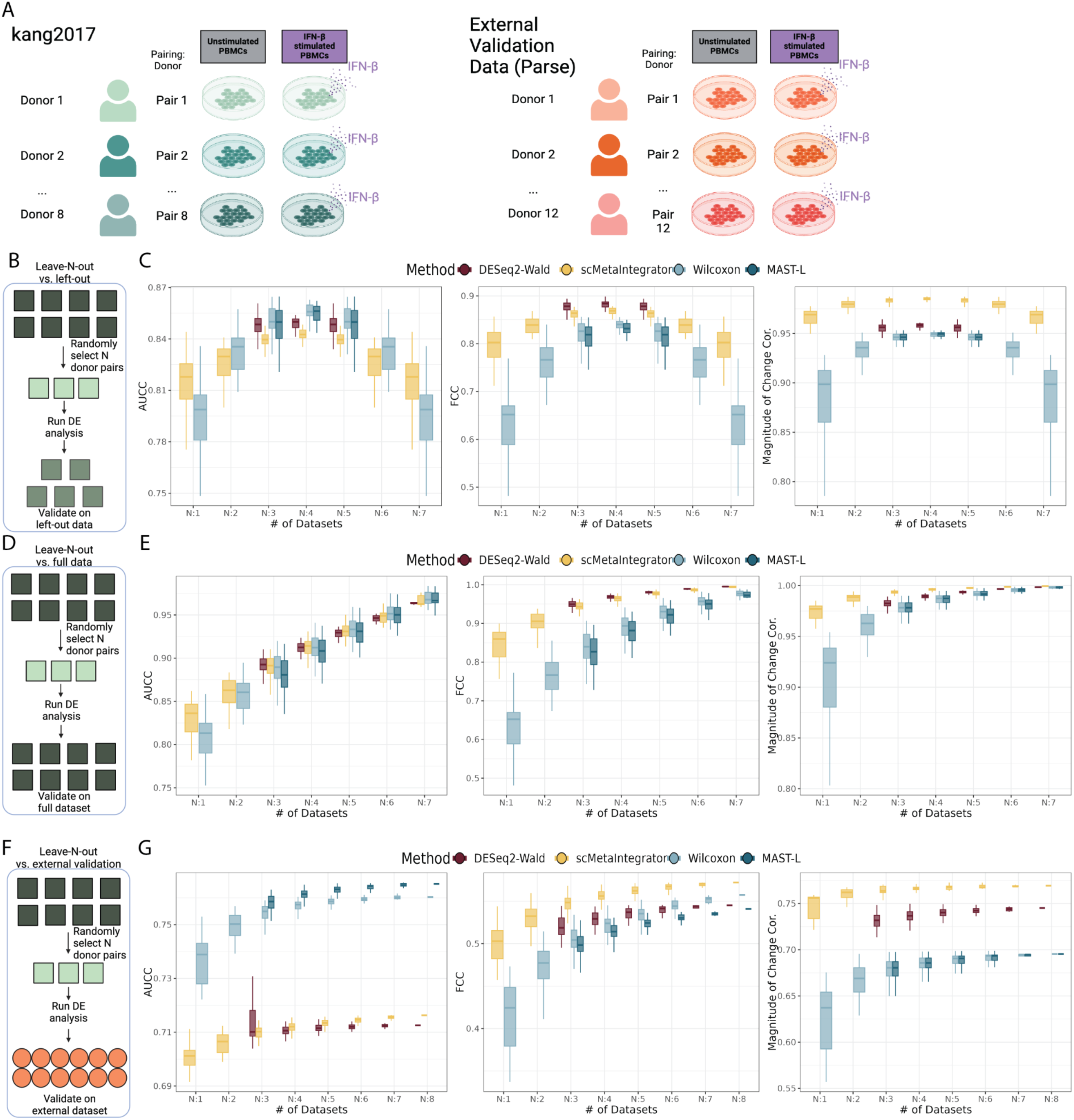
Cross validation of DEGs by method in interferon-beta (IFN-ꞵ) stimulated PBMC datasets. **(A)** Schematics of the test dataset (right) and the validation dataset (left). **(B, C)** Schematic and results of the experiment comparing leave-n-out analyses to the corresponding unseen left-out data analysis. **(D, E)** Schematic and results of the experiment comparing leave-n-out analyses to analyses using the full dataset. **(F, G)** Schematic and results of the experiment comparing leave-n-out analyses to analyses to the results of analyses on separate, validation data. **(C,E,G)** Area under the concordance curve (AUCC) comparison of adjusted p-values on the left. In the middle shows the fold-change correlation (FCC) which is the Spearman’s rank correlation coefficient of the adjusted p-values signed by the direction of change of the magnitude of change values. On the right is the correlation of magnitude of change values.

First, we utilized leave-N-out cross-validation methods to determine within-dataset reproducibility of each method in the held-out samples (**Fig. 5B**). This is also a measure of how well the method performs with potential sample outliers. While AUCC results, which focus solely on adjusted p-values, remained comparable across methods, both scMetaIntegrator and DESeq2-Wald performed better than single-cell methods in analyses that included change in magnitude, which highlighted their robustness to outliers (**Fig. 5C**).

Next, to understand how informative results from a subset of the analysis are in the context of the full dataset, we compared the leave-N-out subset analyses to the results from analysis done across all 8 sample pairs (**Fig. 5D**). While performance solely based on adjusted p-values was similar across method, scMetaIntegrator and DESeq2-Wald outperformed the single-cell methods when considering magnitude of change. Specifically, scMetaIntegrator and DESeq2-Wald analysis on any subset of 3 pairs yielded estimates that were similar to those generated on the full dataset (FCC > 0.9; magnitude of change correlation coef. > 0.96; **Fig. 5E**). These data highlight that despite limited sample sizes, both scMetaIntegrator and DESeq2-Wald provided robust results.

Next, we investigated whether the differences in significance and magnitude of change estimates could impact the functional interpretation of results. Because the cells were specifically stimulated with IFN-ꞵ, we performed pathway analysis to test whether the resulting DEGs from each method could identify the Gene Ontology “response to interferon-beta” gene set. We performed over-representation analysis using over-expressed DEGs (padj < 0.05, magnitude of change > 0) from each method across all leave-N-out analyses and found that all conditions identified the pathway as significantly enriched. Gene set enrichment analysis (GSEA), which considers order of genes used for FCC calculations, demonstrated that while single-cell methods could identify the pathway across all subsets of 3 datasets, scMetaIntegrator identified the gene set as significant (padj < 0.05) with fewer datasets than DESeq2-Wald (**Fig. S4C**). Importantly, normalized enrichment scores for the gene set were consistently higher for scMetaIntegrator than all other methods (**Fig. S4D**). Collectively, these results further highlight that scMetaIntegrator better identifies genes associated with biological changes, when considering statistical significance and effect size estimates, than existing methods.

To determine the reproducibility of each method’s results, we compared each leave-N-out result to those generated when analyzing the complete Parse validation dataset (**Fig. 5F**). Because both our test and validation datasets used IFN-ꞵ stimulation of PBMCs, we hypothesized that results should be comparable across both datasets. There was a total of 5,306 dataset-overlapping genes used for comparisons. When comparing adjusted p-values, Wilcoxon and MAST-L had higher AUCC values than DESeq2-Wald and scMetaIntegrator (**Fig. 5G**). However, when magnitude of change in expression was included in the comparison, scMetaIntegrator has higher FCC and correlation of expression changes (**Fig. 5G**). Together, these data demonstrate that DEGs identified by scMetaIntegrator are stable across both p-value and effect size estimates, producing more reproducible and robust results compared to the other methods.

### Runtime analysis

We compared the speed of scMetaIntegrator to that of other methods by measuring the time required for each method to perform DE analysis by varying the number of cells or sample pairs. We utilized the 29,064 cell kang2017 dataset for this analysis [21].

First, we subsampled 2.5% to 100% of the 29,064 cells in the dataset and randomly distributed them evenly across 6 samples in 3 pairs. Across all subsamples, pseudobulk methods and Wilcoxon rank-sum were fastest, but scMetaIntegrator was significantly faster for larger datasets as compared to all other single-cell methods - including MAST (**Fig. 6A**).

**Figure 6.**
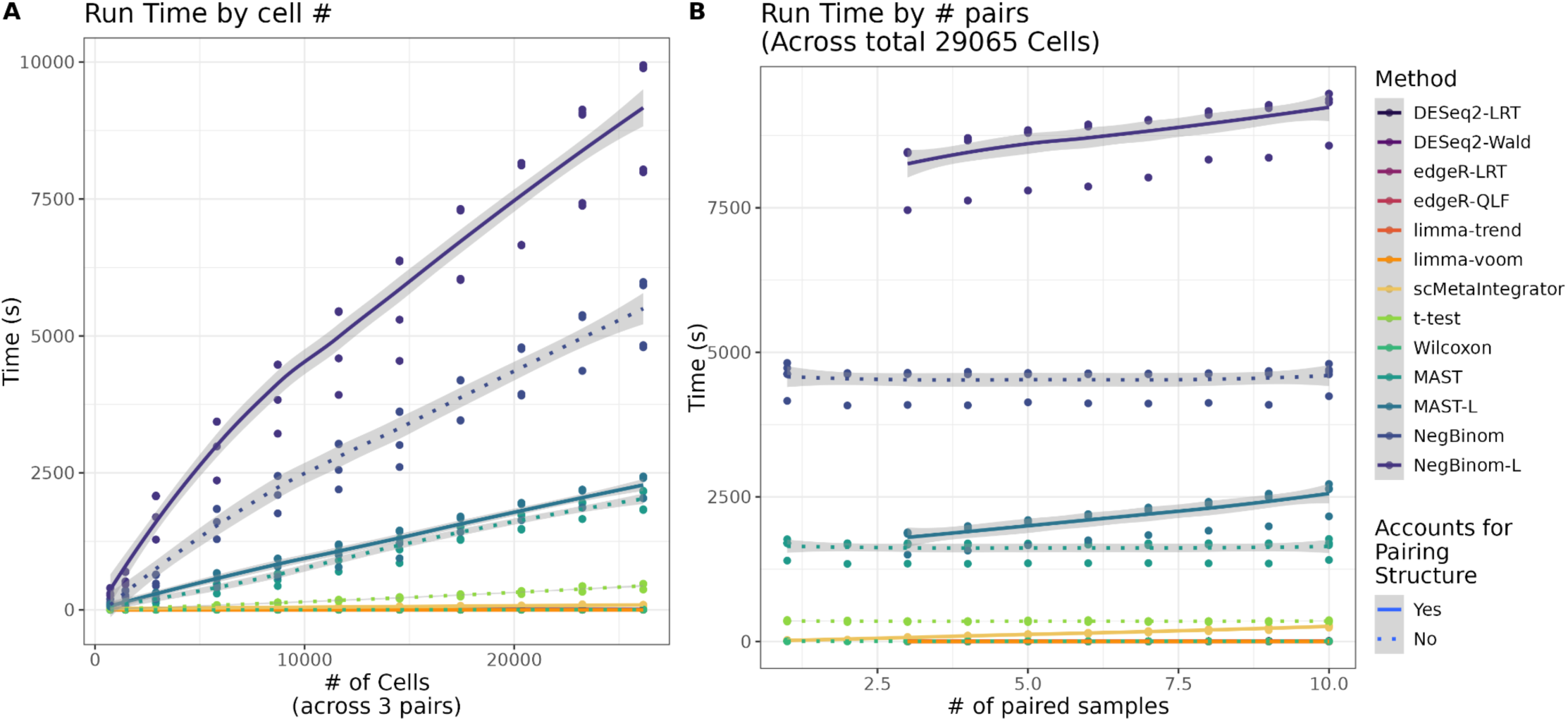
Runtime of each method on scRNA-seq data. **(A)** Time in seconds to run each method on *n* cells evenly divided across 3 pairs. **(B)** Time in seconds to run each method on 29,064 cells evenly split across *n* pairs. 10,374 features were tested for DE across all analyses. Not included in any of the time estimates for pseudobulk methods was the time it took to pseudobulk aggregate where for the maximum combination of **(A)** 29,064 cells across 3 pairs was 0.553 seconds and **(B)** 68,870 cells across 10 pairs was 1.799 seconds.

To test how these methods changed with the number of sample pairs, we randomly divided the 29,064 cells into 1-10 sample pairs. Wilcoxon rank-sum, pseudobulk methods, and t-test demonstrated little difference in time as the number of pairs increased, whereas other methods that account for pairing showed increasing performance speed (**Fig. 6B**). Once again, scMetaIntegrator was significantly faster than the other single-cell methods that did account for pairing structure, highlighting the enhanced usability of this method for paired single-cell data.

## Discussion

Here we present scMetaIntegrator, a differential expression analysis method designed for paired single-cell analysis, which accounts for biological replicate, while also accounting for pairing structure, when generating p-value and effect size estimates unlike existing single-cell methods. Unlike pseudobulk DE methods, scMetaIntegrator also accounts for variation in gene expression across cells within samples and across pairs.

One of the advantages of scMetaIntegrator is its ability to identify conserved changes across paired samples, thereby minimizing the influence of outliers. Our findings demonstrate that scMetaIntegrator identifies DEGs that consistently change across conditions, reinforcing its robustness. We argue that implementing gene thresholding across pairs—rather than solely across conditions—is essential for identifying genes that are associated with the condition of interest, rather than being artifacts of pair-specific variability. To facilitate this, we have integrated our thresholding methodology into the scMetaIntegrator package, enabling its application as a preprocessing step for any differential gene expression analysis.

Moreover, scMetaIntegrator estimated the magnitude and direction of change in DE analyses with higher accuracy and reproducibility than the existing methods. This is a critical aspect, as understanding the directionality of gene expression changes provides deeper insights into the biological implications of the observed changes. While DEG identification is a foundational step in the analytical workflow, the subsequent interpretation of these changes—particularly in the context of the condition of interest—relies heavily on the consistency of directional changes. Many current approaches categorize DEGs into upregulated or downregulated groups, or analyze them based on their order of change in pathway analyses [23]. However, if the directional changes are inconsistent and influenced by outlier samples, the reliability of these genes as true positives is compromised, potentially leading to erroneous biological conclusions.

Our method has several limitations. First, scMetaIntegrator is the requirement of complete pairs. Second, it has only been validated in the context of scRNA-seq; however, we believe that its principles can be extended to other single-cell modalities, such as single-cell ATAC-seq (scATAC-seq) and cytometry by time of flight (CyTOF). Ongoing future expansion of scMetaIntegrator will address both of these limitations.

## Conclusions

Here we present scMetaIntegrator (scMetaIntegrator), an extension of our previously published MetaIntegrator framework to utilize the meta-analysis for performing differential gene expression analysis on paired scRNA-seq data. scMetaIntegrator is an improvement to many single-cell and pseudobulk methods for single-cell differential gene expression by accounting for sample and pairing structures thereby identifying consistently changing genes across analysis pairs. Our method provides robust and reproducible magnitude of change estimates that are resistant to outliers and demonstrates balanced power and false discovery rate estimates between single-cell and pseudobulk methods. In summary, scMetaIntegrator is a useful addition to differential gene expression analysis methods and builds on the approachable MetaIntegrator package and its suite of analysis and visualization techniques.

## Methods

### Data processing of published single cell

For each dataset, pre-processed cell count matrices were used. Generally, we imported the counts and the metadata into Seurat v5.0.2 and normalized the data using the default *Seurat::NormalizeData()* function [24]. Further data-specific details are below:

**Ramírez-Sánchez, et al., 2022.** Metadata and count data was downloaded from the paper’s supplementary files [14]. We only used data from the first timepoint collected and only kept samples with complete pairs. DE analysis was performed by pairing on the assigned matches, and groups used for comparison were the celiac disease subjects and the matched control subjects.

**Arunachalam, et al., 2021.** Metadata and count data was downloaded from GEO (GSE171964) [16]. We only used data from the timepoints 21 and 22 post-vaccination. DE analysis was performed by pairing on the assigned matches, and groups used for comparison. We removed donor 2055 from further analysis due to >99% of a PBMC sample being labeled as monocytes by the authors. DE analysis was performed by pairing on donor, and groups used for comparison were Day 21 and Day 22 samples.

**Kang, et al., 2017.** Metadata and count data was downloaded from GEO (GSE96583) [21]. DE analysis was performed by pairing on donor, and groups used for comparison were IFN-ꞵ stimulated and unstimulated samples.

**Parse Biosciences Dataset.** The “10 Million Human PBMCs in a Single Experiment” dataset was downloaded directly from the Parse Biosciences website as a .h5ad file [22]. Scanpy was used to subset only the unsimulated and IFN-ꞵ stimulated samples. This subset was imported as count and metadata into Seurat. DE analysis was performed by pairing on donor, and groups used for comparison were IFN-ꞵ stimulated and unstimulated samples.

### Generating simulation data

Simulation data was generated using the R package ‘Splatter’ [19,20]. Simulation parameters were built upon the full Kang et al. 2017 dataset using ‘splatEstimate.’

We set up the default cell population to represent having four donors, split into two groups, using the following *splatPopSimulate()* parameters: similarity.scale = 1, batchCells = 10000, 4 samples (donors), similarity.scale = 4, de.prob = 0.2, de.facLoc = 0.1, de.facScale = 0.4, group.prob = c(.5, .5), and eqtl.n = 0.

For outlier sample variation, we added the following to the *splatPopSimulate()* default parameters: condition.prob = c(1/4, 2/4), eqtl.condition.specific = 0.1, cde.facLoc = 0.5, cde.facScale = 0.4. This allowed one donor to have a shifted gene expression profile, independent of the group. To test how varying the outlier donor affected the overall DE results, we generated a Seurat object with the full data. We split the outlier donor and randomly subsetted the object to vary the cells per sample from 50 cells to 5000 cells. For the other three donors, we created a separate object and randomly subsetted each sample (donor + condition) to 1000 cells. We separately merged each of the outlier donor’s objects with the other five donors and ran DE using DESeq2-Wald, scMetaIntegrator, Wilcoxon rank-sum test and MAST with latent.vars = “donor.” We compared each DE result to the ground truth generated on the full dataset by splatter.

To vary between-donor similarity, we did not include outlier sample variation, but we varied the *splatPopSimulate()* ‘similarity.scalè parameter between 0.1 and 10 - which represents the scaling factor for the population variance rate parameter where the higher the value, the increased similarity between the donors. For each parameter variation, we compared each DE result to the ground truth generated by Splatter.

### Gene thresholding for DE analysis

For Figure 1, only genes that were expressed in at least 1% of cells per group (celiac or control) were included in DE analysis. For all further analyses using public, real-world data, only genes that were expressed in at least 1% of cells across each set of paired samples. For simulated dataset analysis, all genes were included for analyses.

### Pseudobulk single-cell data

We utilized two different methods of pseudobulk processing. The default method used was pseudobulk by aggregation (summing) of count values. In fewer cases, we explicitly use averaged pseudobulk expression, in which we average normalized count values.

### Differential gene expression analysis - scMetaIntegrator

For analysis, we started with a dataset’s full Seurat object after normalization. We subsetted the object by each pair grouping, such that each object had two paired samples. The normalized RNA data and associated metadata table for each data subset were imported into a MetaIntegrator object as its own study using the ‘MetaIntegrator’ R package. The class variable of the MetaIntegrator object was set to the Condition/Group comparisons of interest. The final MetaIntegrator object at this point is a list of MetaIntegrator dataset objects for each pair. Then, the MetaIntegrator function ‘runMetaAnalysis’ was performed to run DE analysis. The package to use this method as described is available here: https://github.com/Khatri-Lab/scMetaIntegrator.

### Differential gene expression analysis - single-cell

For Wilcoxon signed-rank test, t-test, negative binomial and MAST methods, we used the Seurat implementation of ‘FindMarkers’ with both ‘min.pct’, and ‘logfc.threshold’ set to 0 and the ‘features’ argument set to the thresholded genes. For the “-L” versions of the negative binomial and MAST method implementations, we included the ‘latent.vars’ argument as the metadata column that samples were paired on (ie. donor for repeat-sampling, pairing column for matched data). The logFC values for each of these methods were the ones generated by Seurat, calculated as the log of the mean of a gene’s expression across the condition cells over the mean of the gene’s expression across the control cells.

### Differential gene expression analysis - pseudobulk

Aggregated pseudobulk data was used as input into pseudobulk DE methods. For all these methods, the design variable incorporated the condition of interest and also the pairing variable. DESeq2 was used with both the Wald test (DESeq2-Wald) and the likelihood ratio test (DESeq2-LRT) [25]. edgeR was used with both the likelihood ratio test (edgeR-LRT) and the quasi-likelihood F-test (edgeR-QLF) [26]. Limma was used with both the trend (limma-trend) and the voom (limma-voom) methods [27].

### Measuring similarity between DE methods

In addition to using Pearson and Spearman’s rank correlation and Jaccard index, we also measured two other metrics based upon Squair, et al. [4]. These metrics are the area under the concordance curve (AUCC) and fold-change concordance (FCC). To calculate the AUCC, we followed the previously published method, ranking genes by their statistical significance values and computing the curve along the entirety of the genes that were measured. To calculate the FCC, we followed the previously published method, multiplying the adjusted P-values by the sign of the magnitude of change between the conditions/groups for each gene and then calculating the Spearman’s rank correlation over all the genes measured between the comparison groups.

### Assessment of DEG heterogeneity

Tau values were calculated per-gene via the MetaIntegrator package. Standard deviation values were calculated across paired samples first taking the aggregated pseudobulk values per-sample, calculating the logFC per paired sample comparison, and then calculating the standard deviation values per-gene across all pairs.

### Gene set analysis

The database used was the Gene Ontology (GO) Biological Process database and the term of interest was “response to interferon-beta” (GO:0035456) [28]. For over-representation analysis, the ‘STRINGdb’ R package [29] was used to perform the analysis on the significantly upregulated DEGs (padj <0.05) by each method. For gene set enrichment analysis, the ‘fgseà R package [30] was used, keeping GO terms with greater than 15 genes and less than 500 genes. For each DE output, genes were ranked in descending order by the sign of the magnitude of change multiplied by the negative log10 of the p-value.

### Runtime analysis

For the runtime analysis, we usd the Kang, et al., 2017 dataset. For analysis of run time by cell number, we randomly subsampled 2.5% to 100% of cells from the full dataset, and evenly split the cells across six samples in three pairs five different times per cell number. For analysis of run time by number of pairs, we took the full dataset and randomly distributed cells across one to ten sample pairs five different times per pairing condition. We performed aggregated pseudobulk on all data subsets that were not included in the overall method times. We ran DE analysis on all of these data subsets, and recorded the time in seconds that it took for the method to run using the R package ‘tictoc’.

### Visualization

All analyses were run in R version 4.2.2. All data visualization was generated in R using ‘ggplot2’ [31] and ‘ComplexHeatmap’ [32]. Schematics were generated with BioRender.

### Code and data availability

All data used are publicly available. Code to replicate analysis will be available here upon publication: https://github.com/Khatri-Lab/scMetaIntegrator_paper. Code for scMetaIntegrator is available here: https://github.com/Khatri-Lab/scMetaIntegrator.

## Abbreviations

padj: adjusted p-value
coef.: coefficient
DE: differential expression
DEG: differentially expressed gene
GSEA: gene set enrichment analysis
logFC: log fold-change
PBMCs: peripheral blood mononuclear cells
SC: single-cell
scMetaIntegrator: single-cell MetaIntegrator
scRNA-seq: single-cell ribonucleic acid sequencing
Wilcoxon: Wilcoxon rank-sum

## Declarations

### Ethics approval and consent to participate

Not applicable.

### Consent for publication

Not applicable.

### Authors’ contributions

KR, CAB, and PK conceived the study and developed the method. CAB and PK supervised the study. KR collected, annotated and processed the data. KR and SNM analyzed the data. KR, CAB, and PK interpreted analysis results and wrote the manuscript. All authors read and approved the final manuscript.

## Acknowledgements

We are grateful to all the labs that conducted the experiments and publicly shared their data, packages and methods. We also thank all study participants and their families for being involved in all included studies. Finally, we thank the members of the Blish Lab and Khatri Lab for their incredibly helpful suggestions throughout the process.

## Author Information

Corresponding author Correspondence to Purvesh Khatri.

## Supplemental figures

**Supplemental Figure 1:**
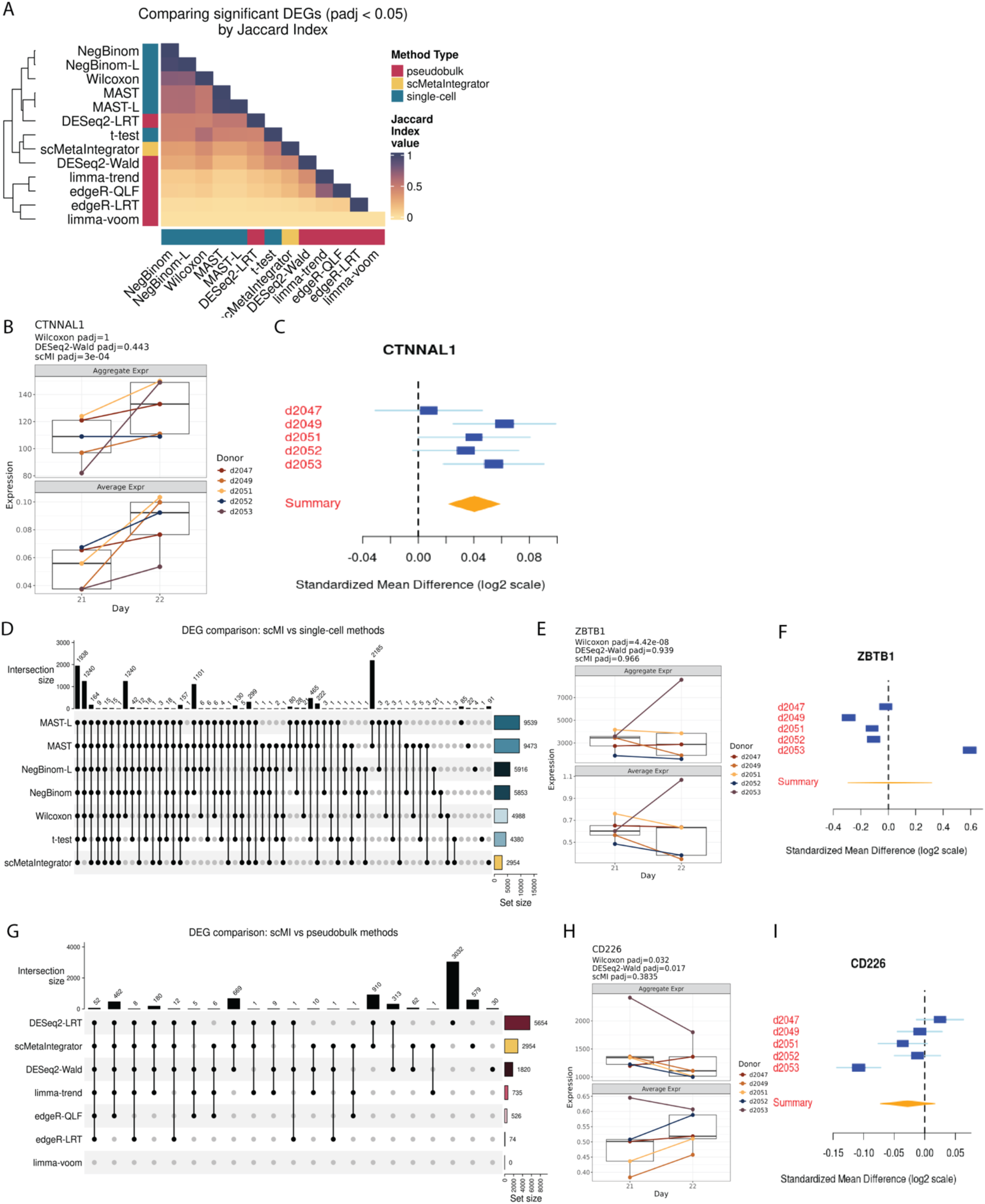
Overlap of DEGs by method. **A)** Jaccard index coefficient of identified DEGs (padj < 0.05) across each method combination. **(B-C)** Focusing on the most significant scMetaIntegrator-unique gene from the 37 genes identified in Figure 3F, **B)** plots of the expression changes by pseudobulk aggregation and average expression methods and **C)** forest plot of the gene across paired single-cell expression. **D)** Upset plot comparing DEGs (padj <0.05) of all single-cell methods and scMetaIntegrator. **(E-F)** Taking the most significant gene by Wilcoxon rank-sum from the 1,240 genes identified across every single-cell method except scMetaIntegrator, **E)** plots of the expression changes by pseudobulk aggregation and average expression methods and **F)** forest plots of the gene across paired single-cell expression. **G)** Upset plot of all pseudobulk methods and scMetaIntegrator. **(H-I)** Taking the most significant gene by DESeq2-Wald of the 5 genes expressed across a majority of methods except scMetaIntegrator, **H)** plots of the expression changes by pseudobulk aggregation and average expression methods and **I)** forest plots of the gene across paired single cell expression.

**Supplemental Figure 2:**
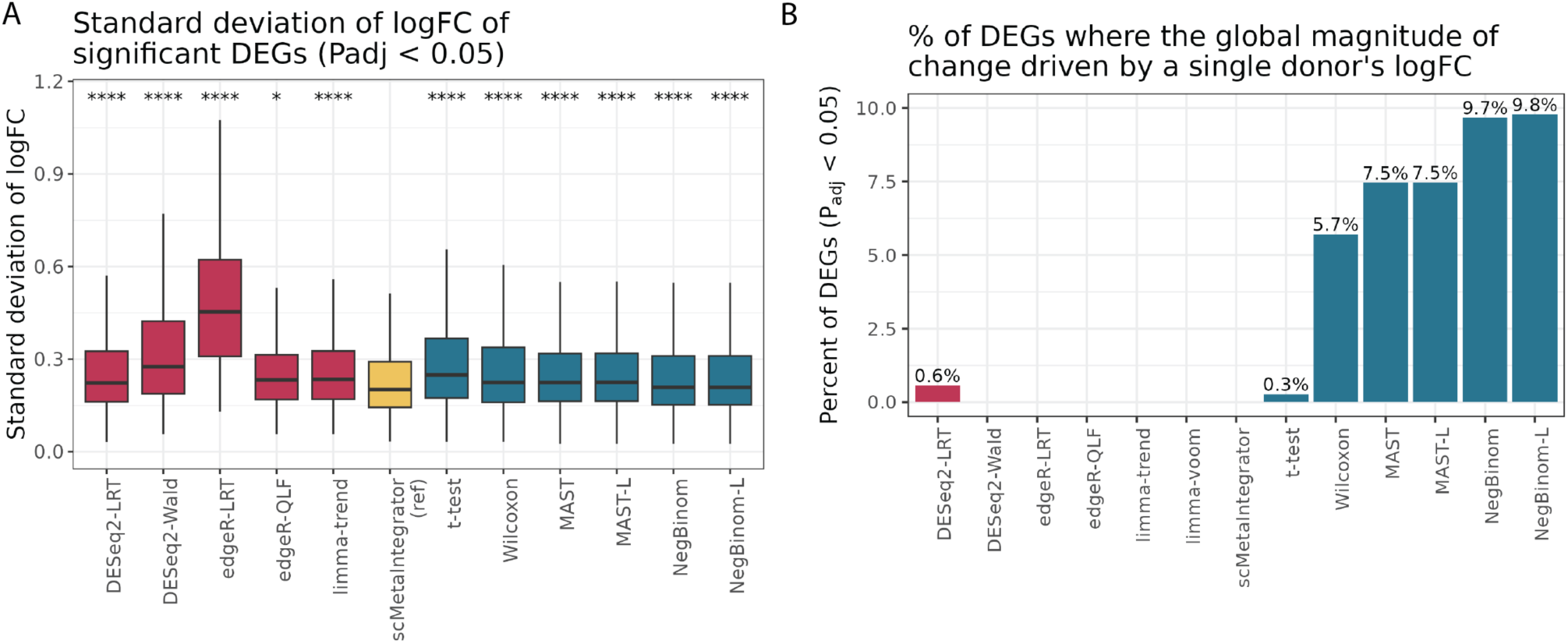
Heterogeneity in Magnitude of Change Value by Method. **A)** Each donor’s logFC was calculated across all genes from their aggregated pseudobulk values. Standard deviation (sd) was calculated per-gene across all 5 paired samples. SD values are plotted for all significant DEGs (padj < 0.05) by each method. Bonferonni adjusted one-sided t-test was performed, testing whether each method had sd values greater than those from scMetaIntegrator. **B)** Barplot of the percentage of each method’s significant DEGs (padj < 0.05) where only one of the five pairs demonstrated a logFC in the direction consistent with method’s magnitude of change value. Asterisk values across figures are represented as follows: ∗padj value < 0.05, ∗∗∗padj value < 0.01, ∗∗∗∗padj value < 0.001, and ∗∗∗∗∗padj value < 0.0001.

**Supplemental Figure 3:**
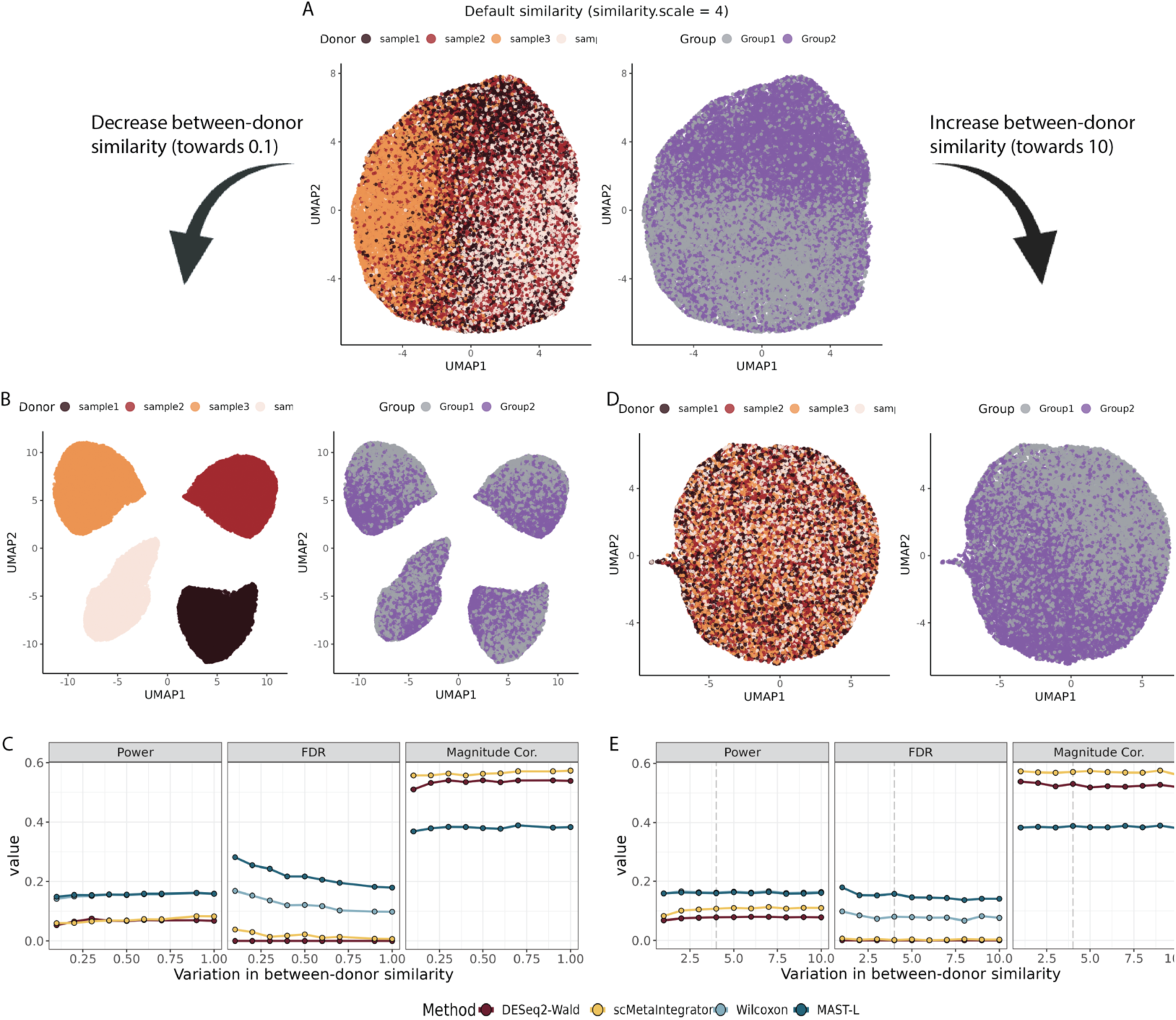
Comparative DE analysis across simulated single-cell data. **A)** UMAPs of the samples when default parameters of simulated data with the splatter similarity.scale parameter set to 4. **B)** UMAPs of the samples when default parameters of simulated data are set to high between-donor variability (similarity.scale parameter set to 0.1). **C)** Comparative analysis of each DE method’s power, FDR, and Spearman’s rank correlation coefficient of magnitude of change values as compared to the ground truth simulation values as between-donor similarity increases (gets closer to 0.1). **D)** UMAPs of the samples when default parameters of simulated data are set to low between-donor variability (similarity.scale parameter set to 10). **E)** Comparative analysis of each DE method’s power, FDR, and Spearman’s rank correlation coefficient of magnitude of change values as compared to the ground truth simulation values as between-donor similarity reduces (gets closer to 10). Grey dashed line indicates the default threshold used in Figure 4 (similarity.scale = 4).

**Supplemental Figure 4:**
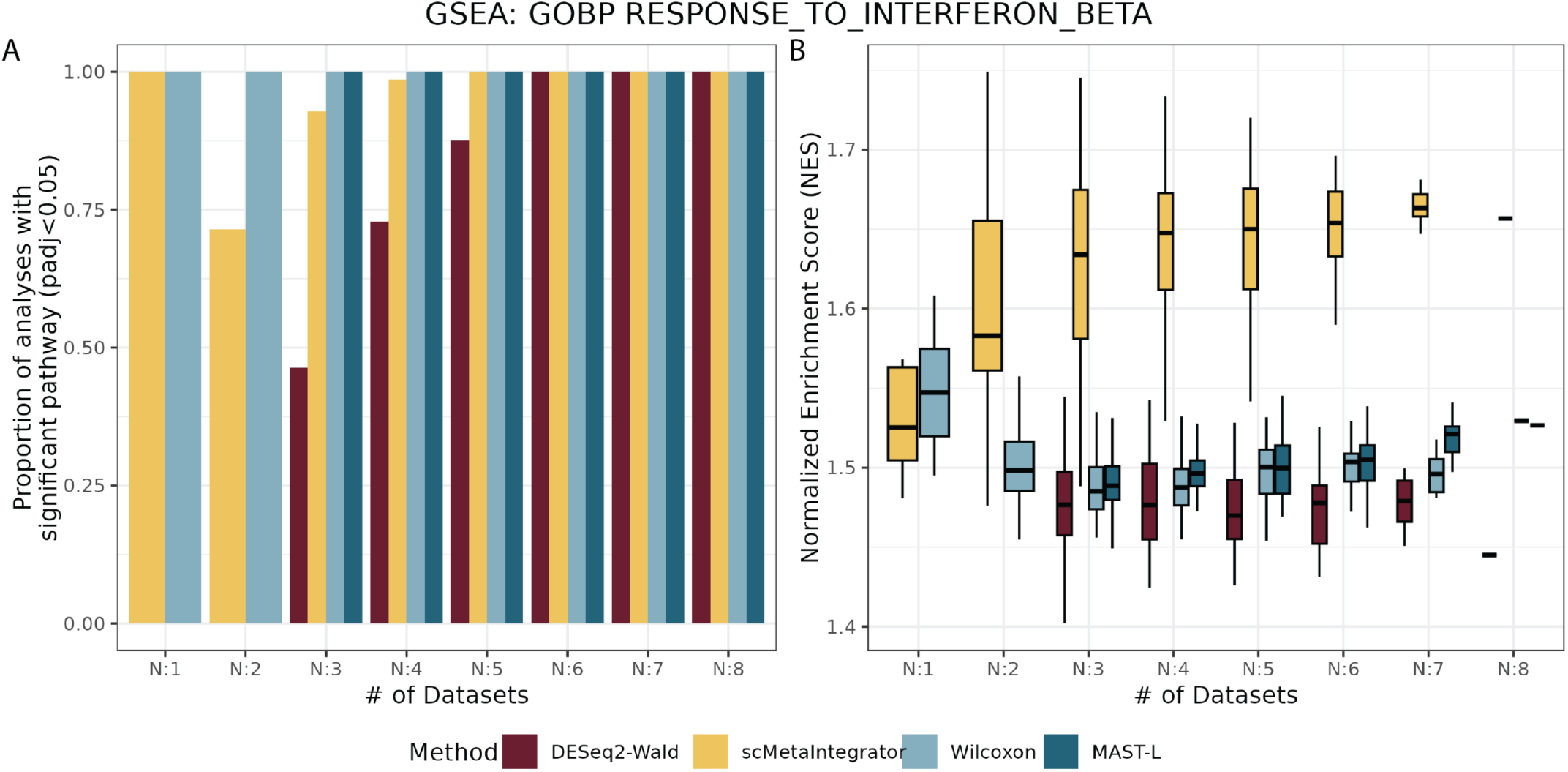
Pathway analysis of cross-validation results. DE analysis was performed across different leave-N-out subsets of the kang2017 dataset, in figure 5A-right. For each analysis, gene set enrichment (GSEA) analysis was performed. **(A)** Proportion plots of the number of analyses in each method and leave-N-out combination that showed the “response to interferon-beta” being significant (padj < 0.05). **(B)** The normalized enrichment score of the “response to interferon-beta” gene set by GSEA analysis in each method and leave-N-out combination, including significant and non-significant results.

